# Uncovering porphyrin accumulation in the tumor microenvironment

**DOI:** 10.1101/2024.04.11.589133

**Authors:** Swamy R. Adapa, Abdus Sami, Pravin Meshram, Gloria C. Ferreira, Rays H.Y. Jiang

## Abstract

Heme, an iron-containing tetrapyrrole, is essential in almost all organisms. Heme biosynthesis needs to be exquisitely regulated particularly given the potential cytotoxicity of protoporphyrin IX, the intermediate preceding heme formation. Here, we report on the porphyrin intermediate accumulation within the tumor microenvironment (TME), which we propose to result from dysregulation of heme biosynthesis concomitant with an enhanced cancer survival dependence on mid-step genes, a process we recently termed ‘*Porphyrin Overdrive’*. Specifically, porphyrins build up in both lung cancer cells and stromal cells in the TME. Within the TME’s stromal cells, evidence supports cancer-associated fibroblasts (CAFs) actively producing porphyrins through an imbalanced pathway. Conversely, normal tissues exhibit no porphyrin accumulation, and CAFs deprived of tumor cease porphyrin overproduction, indicating that both cancer and tumor-stromal porphyrin overproduction is confined to the cancer-specific tissue niche. The clinical relevance of our findings is implied by establishing a correlation between imbalanced porphyrin production and overall poorer survival in more aggressive cancers. These findings illuminate the anomalous porphyrin dynamics specifically within the tumor microenvironment, suggesting a potential target for therapeutic intervention.

## Introduction

Cancer, as a complex and multifaceted spectrum of diseases, remains a significant challenge in medical research and for treatment [1]. The interplay between cancer cells and their microenvironment has revealed novel pathways for understanding and addressing the relentless adversary that is cancer. The molecular landscape of the tumor microenvironment (TME) is diverse and with intricate molecular mechanisms [2, 3]. The emergence of dysregulated metabolism in the TME stands out as a compelling entity with both diagnostic and therapeutic potential [4, 5].

Cancer cells and components of the TME maintain dynamic and mutual interactions to sustain cancer cell survival and proliferation; the tumor cells exhibit distinctive metabolic alterations that contribute to the hallmarks of malignancy [6]. Namely, the Warburg effect, characterized by increased aerobic glycolysis, is a prevalent metabolic shift in which cancer cells favor glycolysis over oxidative phosphorylation even in the presence of oxygen [7, 8]. The heightened dependence on glutamine metabolism supports rapid proliferation, providing essential precursors for biosynthetic pathways and sustaining the tricarboxylic acid (TCA) cycle [9]. Dysregulated lipid metabolism causes increased fatty acid synthesis to fulfill the demands for membrane biogenesis and energy production [10, 11]. One-carbon (1C) metabolism, involving folate-mediated 1C-unit transfers in serine-glycine and methionine syntheses, adjusts to respond to the higher demands for nucleotide synthesis and epigenetic maintenance [12–14]. Redox homeostasis disruption, with elevated reactive oxygen species (ROS) levels, contributes to genomic instability and cancer progression [15]. Furthermore, shifts in amino acid metabolism, such as increased consumption of serine and glycine [14], play pivotal roles in sustaining protein synthesis and other cellular processes.

Porphyrins are cyclic tetrapyrroles with highly conjugated aromatic macrocycles [16]. Because of these π-conjugated structures, porphyrins and porphyrin derivatives exhibit characteristic electronic features, and photophysical and electrochemical properties [17–19], which clearly contribute to their many different biological roles [20]. For example, porphyrin derivatives form chromophoric assemblies involved in light-harvesting for photosynthesis. Metalated porphyrins, typically resulting from chelation of a specific divalent metal ion into the porphyrin macrocyle, have an even larger plethora of functions since the chelated metal ion can bestow redox activity and an additional ligand coordination site. Ferrous iron (Fe^2+^)-bound protoporphyrin IX (PPIX) or heme b (thereon, referred to as heme) stands out as a metalated porphyrin with biochemical and physiological activities that go beyond oxygen transport, oxygen storage, electron transfer and catalysis [20]. Heme-bound proteins act as heme-responsive sensors and regulate transcription [20–24], microRNA splicing and processing [25], translation [26], heme degradation [27], protein degradation [28, 29], K^+^ channel function [30], redox status [30–32] and protein-protein interactions [20]. The enzyme-catalyzed chelation of Fe^2+^ in PPIX corresponds to the terminal step of the heme biosynthetic pathway [33–35]. This reaction is finely regulated as both PPIX and iron ion are potentially cytotoxic [36, 37]. However, and in a somewhat ironic twist, the uncontrolled production and accumulation of PPIX, typically observed in cancer, can serve as both a diagnostic marker and a therapeutic agent [38]. We recently coined the term ‘Porphyrin Overdrive’ to describe this process [38]. The fluorescence emission properties of porphyrins like PPIX enable non-invasive tumor imaging and cancer detection [39]. Because porphyrins are photosensitizers, they are employed in photodynamic therapy (PDT) [40]. Light activation of the porphyrins accumulated in cancer cells leads to ROS generation and their selectively targeted cell death [40].

The TME is a dynamic and heterogeneous milieu of cells where the tumor resides. It comprises normal cells, including immune and stromal cells, signaling molecules, extracellular matrix, and blood vessels that enclose and feed the tumor cells [3]. Recent investigations have unveiled that dysregulation of heme biosynthesis within the TME supports cancer [41, 42]. In fact, metabolic patterns for heme production and transport are altered in comparison to those in normal cells [41–44]. In this study, we delineate a novel facet of metabolic rewiring in the TME. We demonstrate that porphyrin production occurs in both cancer cells and stromal cells in the TME. Porphyrins (*i.e.,* heme intermediates) play a prominent role as their biosynthesis, and not that of heme *per se*, is linked to cancer aggressiveness. Significantly, porphyrin production not only sustains cancer cell survival and proliferation but also influences clinical outcomes. Thus, understanding the dynamics of porphyrin production in the TME holds the promise of unveiling novel insights into the metabolic adaptations that sustain tumorigenesis.

## Materials and Methods

### Cell Culture, Growth, and Quantification

Cell culture, proliferation, and quantification procedures were carried out utilizing human lung cancer cell lines. Cells were cultivated in RPMI 1640 medium (Gibco), supplemented with 10% fetal bovine serum (Sigma), 2 mM L-glutamine, and additional components such as gentamicin (1000x, Fisher, Cat No. 15-710-072) and PennStrepNeo solution (100x, Fisher, Cat No. 15640-055). The incubation of cells occurred at 37 °C in a humidified 5% CO_2_ atmosphere to facilitate optimal growth conditions. The stock solutions of 5-aminolevulinic acid hydrochloride (ALA) and glycine were prepared by dissolving them in distilled water and phenol red-free culture medium, respectively. ALA, sourced from Alfa Aesar (Cat No. A16942ME), was stored at –20 °C with a stock concentration of 1.0 M. Glycine, acquired from Fisher Chemical (Cat No. BP381-500), was stored as a 1 M stock solution. Cancer cells were initially plated at a density of 1 – 2 x 10^5^/mL, and cell counting, performed using trypan blue exclusion (Corning, Cat No. 25-900-CI), was carried out over time. As part of the quantification process, total cell numbers were determined based on the passage dilution at each time point.

### Heme Biosynthetic Pathway-related Gene Expression Analysis

Analysis of the gene expression patterns for the enzymes of the heme biosynthetic pathway in tumors and normal tissues was performed using the Genotype-Tissue Expression (GTEx) project [45, 46] and The Cancer Genome Atlas (TCGA) program [47] as data resources. The heme biosynthetic pathway gene expression patterns in tumors *vs*. matched normal tissues were used for analysis. Briefly, this analysis involved a thorough comparison of gene expression profiles for enzymes involved in the heme biosynthetic pathway between tumor samples and their corresponding paired normal tissues. Non-parametric pairwise Wilcoxon comparisons were conducted between normal and tumor samples. The expression levels of each gene involved in heme biosynthesis in both normal and tumor tissues were examined, and specific changes associated with each biosynthetic step in the linear pathway were analyzed to infer significant expression differences between tumor and normal tissues.

### Heme CRISPR Cancer Gene Essentiality Analysis

The essentiality analysis of cancer genes using CRISPR/Cas9 was conducted utilizing data sourced from the DepMap Portal, following established methodologies [48–50]. Whole-genome CRISPR/Cas9 datasets were employed to identify significantly depleted growth of mutant cells resulting from specific gene knockouts in pooled experiments. Gene essentiality was inferred from the dependency of a given gene, deduced from CRISPR/Cas9 gRNA-mediated gene knockout. Essential scores were utilized to assess cell growth fitness, with lower scores indicating a greater impact on cell viability upon gene loss. Specifically, scores of 0, < 0, and > 0 denoted no fitness change, fitness loss, and fitness gain (implying potential growth advantage for the cell line) under the assay conditions. To address copy number bias in whole-genome CRISPR/Cas screens, the method outlined in [48, 49] was applied, involving the computation of the mean of sgRNAs *versus* the control plasmid library.

Identification of commonly essential genes was based on their significance for the fitness of most cell lines across various cancer types [51, 52]. For the analysis of *in vivo* gene essentiality in pancreatic and lung cancer models, data published in [53] were used to specifically investigate genes related to the heme biosynthetic pathway enzymes and heme transporters.

### *In vitro* Primary Human Cell Culture, Quantification, and Imaging

384-well plates (Greiner, Cat No. 781091) were aseptically handled in a class II biosafety cabinet and then placed in a secondary container, serving as a lid to control evaporation (*i.e*., plates were positioned in large assay pans). A day prior to cell seeding, the wells were collagen-coated with 40 μL of 15 μg μL-1 rat tail collagen I (Corning, Cat No. 354236) in sterile filtered 0.02 M acetic acid (Thermo Fisher Scientific, Cat No.) and maintained at 37 °C overnight. Immediately before seeding, wells were washed three times with sterile phosphate-buffered saline (PBS) and then filled with 20 μL in vitro GRO® CP plate medium (BioIVT, Cat No. Z99029) supplemented with 1x Pen-Strep-Neo solution (100x, Fisher, Cat No. 15640-055) and 20 μM gentamicin (1000x, Fisher, Cat No. 15-710-072). Cryopreserved primary human cells (BioIVT) were thawed by immersion in a 37 °C water bath for 2 minutes, sterilized with 70% ethanol in a sterile field, and their contents were directly added to 4 mL of plate medium. Live and dead cells were quantified through trypan blue exclusion on a Neubauer improved hemocytometer. The cell density was adjusted to 1 × 10^3^ live cells μL^-1^, and 18 μL of cell suspension was added to each well. Medium exchanges with the GRO® CP plating medium, as described earlier, were performed thrice weekly. The cells were incubated with 1.0 mM ALA at 37 °C for 4 hours, and both ALA-treated and non-treated cells were handled under very low light conditions. During the last 45 minutes of incubation, a staining solution, diluted in phenol-free, serum-free RPMI (Gibco, Cat No. 11835055), and containing Hoechst 33342 (Life Technologies, Cat No. H3570) at a final concentration of 10 μM, was added to the cells. Live cell imaging was conducted on a CellInsight CX7 High-Content Screening Platform (Thermo Fisher Scientific), with each well counted for cell nuclei staining.

### *in vitro* Cancer Cell Line Culture, Quantification, and Imaging

Cells were allowed to grow until reaching 70%. In the *in vitro* culture of cancer cell lines, cryopreserved cells were thawed, suspended in a pre-prepared culture medium, and then transferred to a collagen-coated T75 flask (Corning, Cat No. 354236) at a density of 5 μg/cm^2^. The culture medium was composed of a 1:1 (v/v) mixture of F12 base medium (Invitrogen, Cat No. 11765-054) and MEM base medium (Invitrogen, Cat No. A10490-01), supplemented with 10% FBS (Hyclone, Cat No. SH30070), 1.0 M HEPES (Invitrogen, Cat No. 15630-080), and 200 mM glutamine (Invitrogen, Cat No. 25030-081) [54]. Cells were allowed to grow until reaching 70% confluence, with medium changes every other day. Upon reaching the desired confluence, cells were trypsinized using TrypLE™ Express Enzyme (1X) (Gibco, Cat No. 12605028), washed with culture medium, and then seeded at a density of 6000 cells/well in 384-well plates (Greiner, Cat No. 781091). The cells were cultured in 20 μl of the above medium per well and incubated at 37 °C either in the absence or presence of 1.0 mM ALA for 4 hours. Both ALA-treated and non-treated cells were handled under very low light conditions.

During the final 45 minutes of incubation, a staining solution, diluted in phenol-free, serum-free RPMI (Gibco, Cat No. 11835055) and containing Hoechst 33342 (Life Technologies, Cat No. H3570) at a final concentration of 10 μM, was added to the cells. Live cell imaging was conducted using a CellInsight CX7 High-Content Screening Platform (Thermo Fisher Scientific), with each well of the 384-well plate capturing images from 15 fields at 20x magnification for nuclei staining.

### Cellular PPIX Quantification

Quantification of intracellular PPIX accumulation was conducted through fluorescence-activated cell sorting (FACS), following established procedures [55]. Cancer cells were incubated in medium (outlined in the Cell Culture section) containing 1.0 mM ALA at 37 °C for 4 hours, under low light conditions. Post-incubation, cells underwent triple washing with Dulbecco’s phosphate-buffered saline (DPBS, 1X, Ca^2+^– and Mg^2+^-free; Corning, Cat No. 21-031-CV) and were resuspended in 250 μL of 1x DPBS. Subsequently, cells were washed once with serum-free medium (Gibco, Cat No. 11835055) and cultured in 6-well plates with the designated medium (described in the Cell Culture section), with or without the addition of ALA (0.1 mM, 0.25 mM, 0.5 mM, and 1.0 mM) at 37 °C. Intracellular PPIX concentration was assessed 18 hours later using FACS.

The cell suspension of dissociated fibroblasts, pre-treated with a 40-μm Flowmi™ Cell Strainer to exclude clumps and debris, was transferred to BD Falcon tubes under minimal light exposure to mitigate potential phototoxicity from PPIX accumulation.

FACS analyses were executed using a BD LSR II Analyzer (Becton, Dickinson, and Company) alongside FACSDiva Version 6.1.3 software. To eliminate background red fluorescence, the 633 nm-red laser was disabled during PPIX emission data collection.

PPIX emission within the 619 nm and 641 nm range (630/22BP filter) was determined following excitation with the 405 nm laser. Forward-scatter (FSC) versus side-scatter (SSC) dot plots were employed to gate the entire cell population, excluding cell debris. A minimum of 10,000 cells within the gated region were then represented in dot plots of SSC *vs*. PPIX fluorescence, with the gate defined using cells lacking perturbation as negative controls.

### Patient Tumor Gene Expression Analysis and Survival Analysis

We analyzed gene expression data from The Cancer Genome Atlas (TCGA) to investigate the heme biosynthesis process in lung adenocarcinoma and its association with patient survival outcomes. Specifically, we examined the expression levels of ALA dehydratase (ALAD) and hydroxymethylbilane synthase (HMBS), representing the second and third enzymes in the heme biosynthetic pathway, using the TCGA dataset. Kaplan-Meier survival curves [56] and log-rank tests [57], were performed to assess the relationship between gene expression levels and overall survival (OS). Patients were stratified based on high and low gene expression levels, and statistical significance was determined using log-rank tests.

## Results

### Porphyrin Accumulation Is Absent In Normal Cells but Present in Lung Cancer Cells

Metabolic dysregulations play an important role in the complex landscape of lung cancer, contributing to its progression, resistance to treatment, and overall clinical outcomes [28–30]. To evaluate if lung cancer cells, and their TME, over produce porphyrins, we first investigated and compared heme biosynthesis in diverse types of non-cancerous cells, *i.e.,* normal differentiated cells, replicating fibroblasts, human primary cells, and human primary hematopoietic stem cells. We looked for evidence of porphyrin production, defined by a non-homeostatic or imbalanced heme biosynthetic pathway and heightened accumulation of intermediates. ALA, or 5-aminolevulinic acid, is utilized as a precursor to bypass the rate-limiting first step of heme biosynthesis, enabling porphyrin accumulation in an unbalanced pathway; however, no accumulation occurs when the heme biosynthesis pathway is balanced.

We found normal human cells do not over produce porphyrins, a result that is consistent with our recent study [38] and many other prior reports [39, 58]. Here, we experimentally validated that primary human lung fibroblasts lack porphyrin accumulation, *i.e.*, 0% of the tested normal lung cells accumulate PPIX after addition of exogenous ALA (Fig. 1A). By contrast, high-resolution imaging of human lung cancer H460 cells confirms that PPIX accumulates and is distributed throughout the cell – from mitochondria, where PPIX is synthesized, to cytosol (Fig. 1B).

**Fig. 1.**
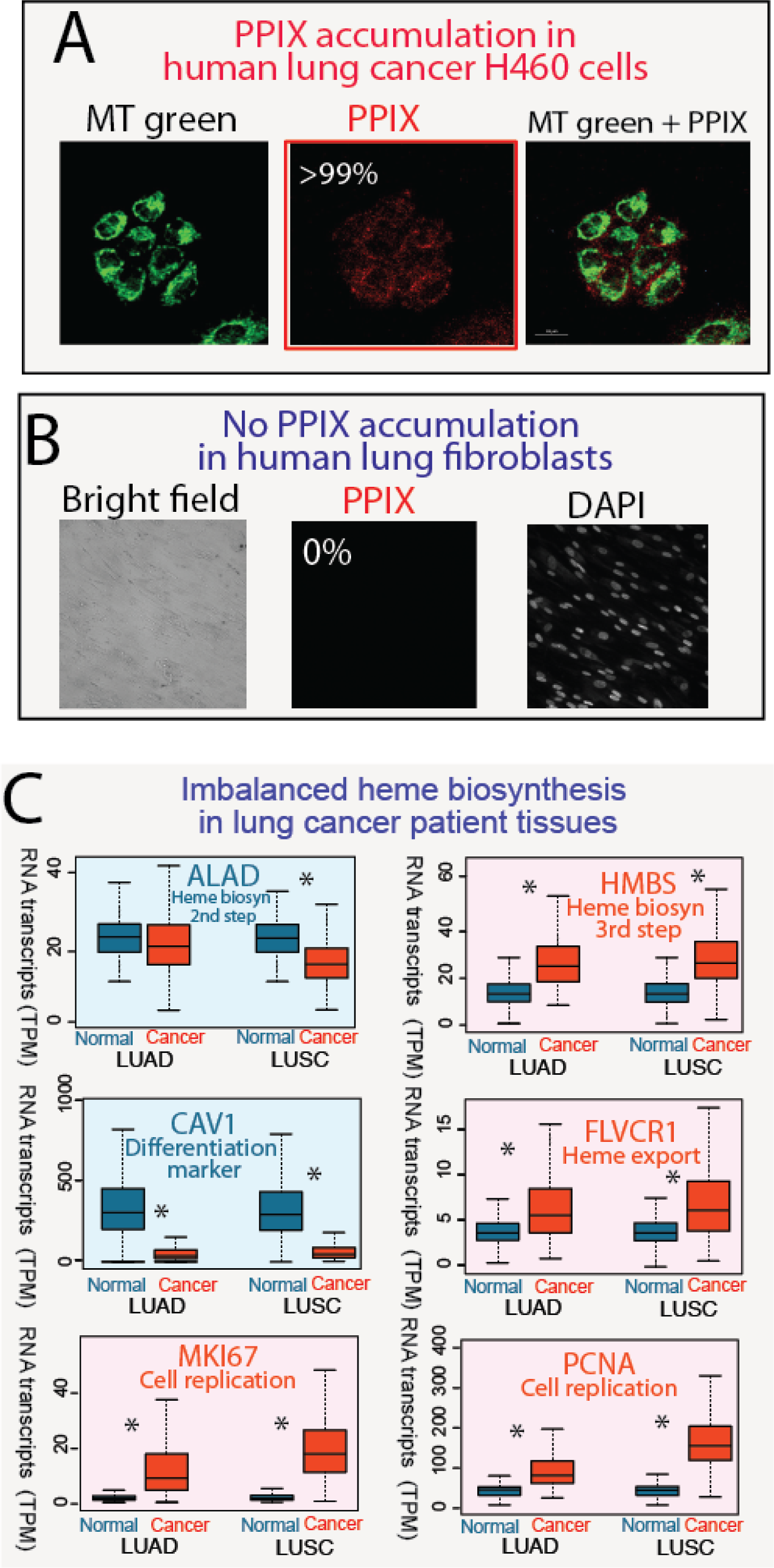
Human lung cancer cells, but not lung fibroblasts, accumulate PPIX. **A.** After treatment of H460 cells with 1.0 mM ALA for 4 h, the confocal fluorescence images were obtained. From left to right, the panels correspond to (1) green fluorescence, (2) red fluorescence, and (3) superimposed green and red fluorescence images. The green arises from the selective labeling of mitochondria with the MitoTracker Green (MT green) probe (excitation wavelength, *λ*_ex_, at 490 nm and emission wavelength, *λ*_em_, at 516 nm), and the red fluorescence is due to PPIX fluorescence (*λ*_ex_ 405 nm; *λ*_em_, 635 nm). > 99%, over 99% of H460 cells accumulate PPIX. The frame for the pertinent image is colored in red. Scale bars represent 20 μm. **B.** Primary human lung fibroblasts do not accumulate PPIX. From left to right, the panels correspond to (1) brightfield, (2) red (PPIX) fluorescence, (3) blue fluorescence. PPIX, which fluoresces red, is not detected (middle panel). 0%, no primary lung fibroblasts accumulate PPIX. The blue fluorescence is due to binding of DAPI to adenine– thymine-rich regions of nuclear DNA (*λ*_ex_ 375 nm; *λ*_em_, 460 nm). **C.** Gene expression patterns from TCGA lung cancer patients indicate imbalanced heme biosynthetic pathway and enhanced heme transport. Lung epithelial cell differentiation (CAV1) and cell proliferation (MKI67 and PCNA related genes are plotted as controls. Statistically significant differences are indicated by *. [Abbreviations: DAPI, 4’,6-diamidino-2-phenylindole; LUAD, lung adenocarcinoma; LUSC, lung squamous cell carcinoma]

Results of our analysis of gene expression data for lung cancers and matching normal tissue are illustrated in Fig. 1C (Supplemental Table S1). While the expression of *ALAD*, encoding the enzyme for the second step of the heme biosynthetic pathway, is either unaffected or downregulated, the expression of *HMBS*, encoding the third enzyme of the pathway, and the expression of *FLVCR1*, coding for the heme transporter 1, are upregulated in patient tumors (adjusted pValue < 0.01). Similarly, the expression of genes for cell differentiation (CAV1 for caveolin-1) [59] and cell proliferation (MKI67 for marker of proliferation Ki-67 and PCNA for proliferating cell nuclear antigen) [60, 61] were down– and upregulated, respectively, in tumor samples (adjusted pValue < 0.01) (Fig. 1C).

### CRISPR analysis shows porphyrin production contributes to tumorigenesis

To study the molecular and genetic mechanisms that underlie the reliance cancer on heme metabolism, we assessed the data from genome-scale CRISPR/Cas9 loss-of-function screens of human lung cancer cell lines. Towards our central focus on the gene dependencies related to the eight enzymatic steps of the heme biosynthetic pathway, we calculated the respective gene essentiality scores, akin to the lethality scores utilized in constructing a cancer dependency map (DepMap)[48–52]. Figure 2 illustrates the survival dependence of 19 metastatic lung cancer cell lines and 28 primary lung cancer cell lines on heme biosynthesis and trafficking. Gene essentiality score values below zero signify a reduction in cell growth/viability upon the loss of gene X. Therefore, the lower the essentiality score of gene X, the greater the dependence of cells on that specific gene (Supplemental Table S2).

**Fig. 2.**
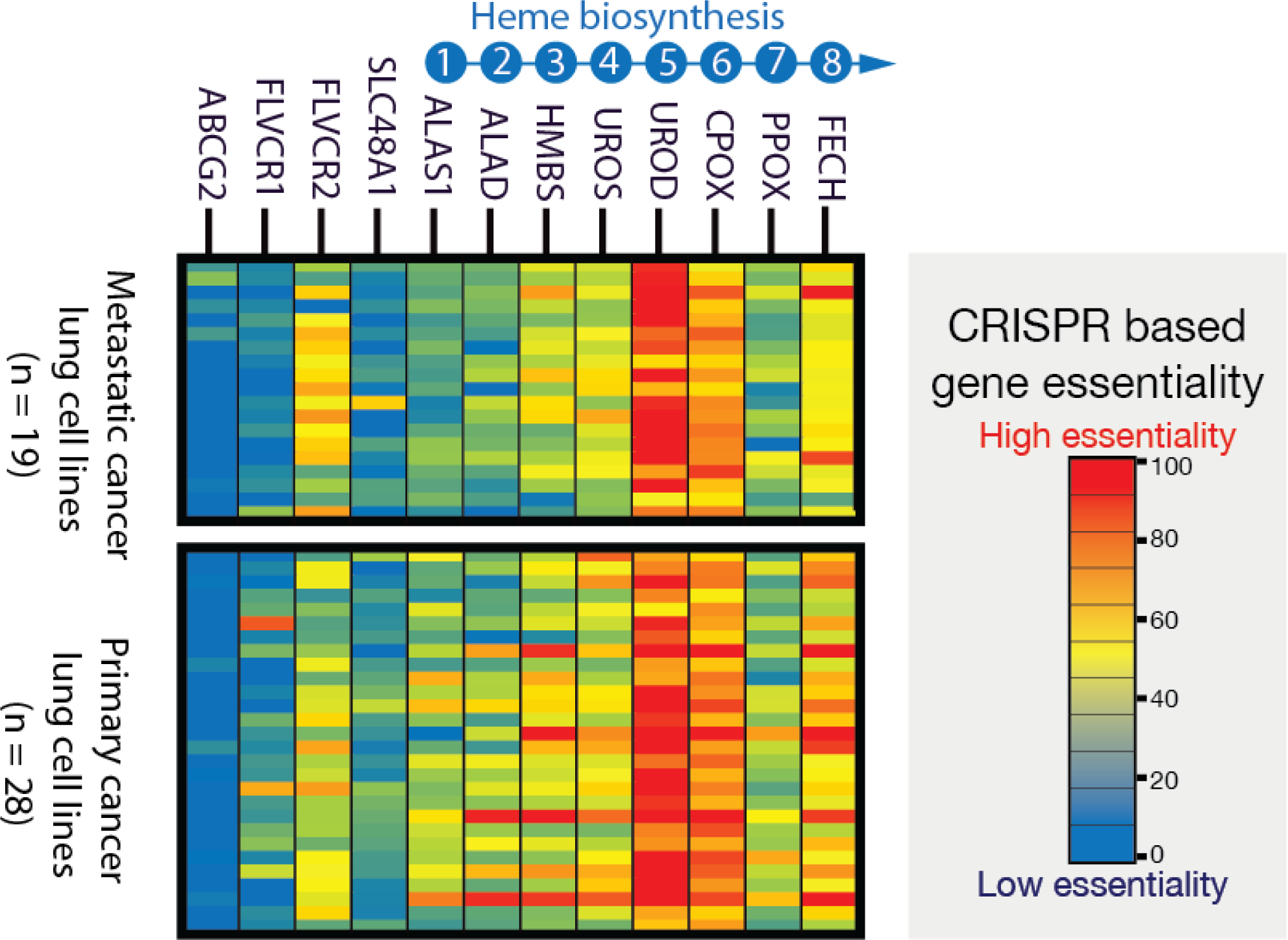
CRISPR/Cas9 gene targeting and inferred gene essentiality in lung cancer cells show porphyrin production. Whole genome loss-of-function growth phenotypes show that a partially functional heme biosynthetic pathway to survive. Gene essentiality is estimated from gene X dependency inferred from CRISPR/Cas9 gRNA gene X knockout. Low and high cell viability with deletion of gene X indicate that the cancer cells have a high and low dependence on gene X for survival, respectively. Thus, the lowest and highest gene essentiality values are associated with the least and most profound dependence of the cells on the loss of gene X, respectively. The UROD gene for the fifth enzyme of the pathway has the highest essentiality in cancers, while genes encoding other enzymes are dispensable in many cancer cell lines. Each of the rows represents a distinct cancer cell line derived from primary and metastatic tumors. Numbers 1 to 8 indicate the order of the eight enzymes of the heme biosynthetic pathway. [ABCG2, ATP-binding cassette (ABC) transporter subfamily G, member 2; ALAD, ALA dehydratase (aka porphobilinogen synthase); ALAS1, 5-aminolevulinate synthase 1; CPOX, coproporphyrinogen oxidase; FECH, ferrochelatase; FLVCR, feline leukemia virus subgroup C receptor family; HMBS, hydroxymethylbilane synthase; PPOX, protoporphyrinogen oxidase; SLC48A1, solute carrier family 48, member 1, a.k.a heme transporter HRG1; UROD, uroporphyrinogen III decarboxylase; UROS, uroporphyrinogen III synthase]

Remarkably, instead of encountering the expected fully balanced heme biosynthetic pathway characteristic of normal cells—where all steps exhibit similar essentiality—we observed a cancer cell dependency on various pathway steps in a “non-balanced” manner (Fig. 2). Significant genetic essentiality differences were found among genes encoding each step of the pathway (Supplemental Table S3). Specifically, the first step ALAS1 showed no concordance with mid-step genes in essentiality. Notably, the highest cancer gene essentiality was attributed to the genes encoding enzymes responsible for the fifth and sixth steps of the pathway, namely uroporphyrinogen III decarboxylase (UROD) and coproporphyrinogen III oxidase (CPOX), respectively. This lung cancer cell line outcome was consistent with our recent study [38]. These findings reveal the “unbalanced” nature of heme biosynthesis in cancer cells, wherein many cancer cells can survive without functional first and terminal enzymatic steps but demonstrate dependence on the intermediate steps.

Furthermore, the gene essentiality analysis emphasized the significance of heme trafficking, exemplified by the heme importer FLVCR2, in cancer cells for at least *in vitro* survival (Fig. 2). To investigate cancer heme metabolism *in vivo*, we analyzed previously published in vivo mouse CRISPR/CAS9 loss-of-function and metabolic essentiality data obtained from lung cancer models [53]. The heightened in vivo essentiality of mid-step heme biosynthesis genes (HMBS, UROS, CPOX, and PPOX) in the examined lung cancer models, as reported in the aforementioned dataset, suggests a notable dependency on heme intermediates, particularly porphyrin production. Contrary to the authors original interpretation, which suggested enhanced in vivo heme biosynthesis, our step-by-step analysis[38] indicates a likely enhancement of intermediate production, as neither the first nor the last step genes are essential *in viv*o in lung cancer [38]. Notably, we highlight an increased genetic reliance on these mid-step genes for cancer survival, observed during the transition from *in vitro* to *in vivo* conditions. This observation indicates an intensified requirement for the unbalanced pathway of porphyrin accumulation within the tumor tissue environment.

### Porphyrin Production Occurs Primary Tumor Cancer-associated Fibroblasts (CAFs) Residing in TME

Remarkably, cancer cells, both *in vitro* (Figs. 2) and *in vivo* [53], do not necessitate endogenous heme synthesis based on genetic inferences. However, the genes encoding hemoproteins (*i.e.,* proteins requiring heme as a cofactor) such as cytochrome C is characterized as top 10% most essential genes [38] in the largest collection of cancer lines in DepMap (version 23Q4), and the heme/intermediates transport gene FLVCR proves to be required (Fig 2). This discrepancy between the “defect” in endogenous heme production and the necessity for heme-cofactors within the cells, coupled with the overproduction of porphyrins, indicates the essential nature of trafficking activities. This observation suggests the existence of a specialized cancer microenvironment tailored for efficient metabolite trafficking. Operating within a complex network, cancer cells exhibit a suggested heme flux associated with porphyrin production, characterized by heightened heme and intermediates trafficking. This notion is supported by the genetic dependence on heme trafficking genes (Fig. 2) and the elevated expression of trafficking genes in patient tumors (Fig. 1C). study the molecular and genetic mechanisms that underlie cancer’s reliance on heme upregulated, respectively, in tumor samples (adjusted pValue < 0.01) (Fig. 1C).

Nevertheless, the source and fate of trafficked heme/porphyrins in the TME remain unclear at present. While heme trafficking is tightly regulated in the normal marrow environment due to potential heme toxicity [55, 62, 63], the relationship between porphyrin production and the specific cancer microenvironment optimized for heme transport and detoxification remains to be fully elucidated. Of particular interest is the role played by cancer-associated fibroblasts (CAFs), a cell population recognized for its significant contributions to shaping the TME [64]. To study cancer cells and TME at single cell resolution, we used single-cell lung cancer transcriptomes generated from Zilionis et al. [65] and analyzed heme biosynthesis and trafficking in diverse cell populations present in the tumor (Supplemental Table S4). HMBS and FLVCR1 were the most upregulated heme biosynthesis genes in tumor cells from 3 different NSCLC lung cancer patients (Fig. 3) (Supplemental Table S4). These single cell transcriptomes indicate that cancer cells exhibit features of porphyrin production, consistent with the abnormal heme anabolism and endogenous PPIX accumulation upon ALA administration in cancer cells compared to normal cells (Fig. 1C). By contrast, major groups of stromal and immune cell populations do not show evidence of imbalanced heme biosynthesis or heme trafficking (Fig. 3).

**Fig. 3.**
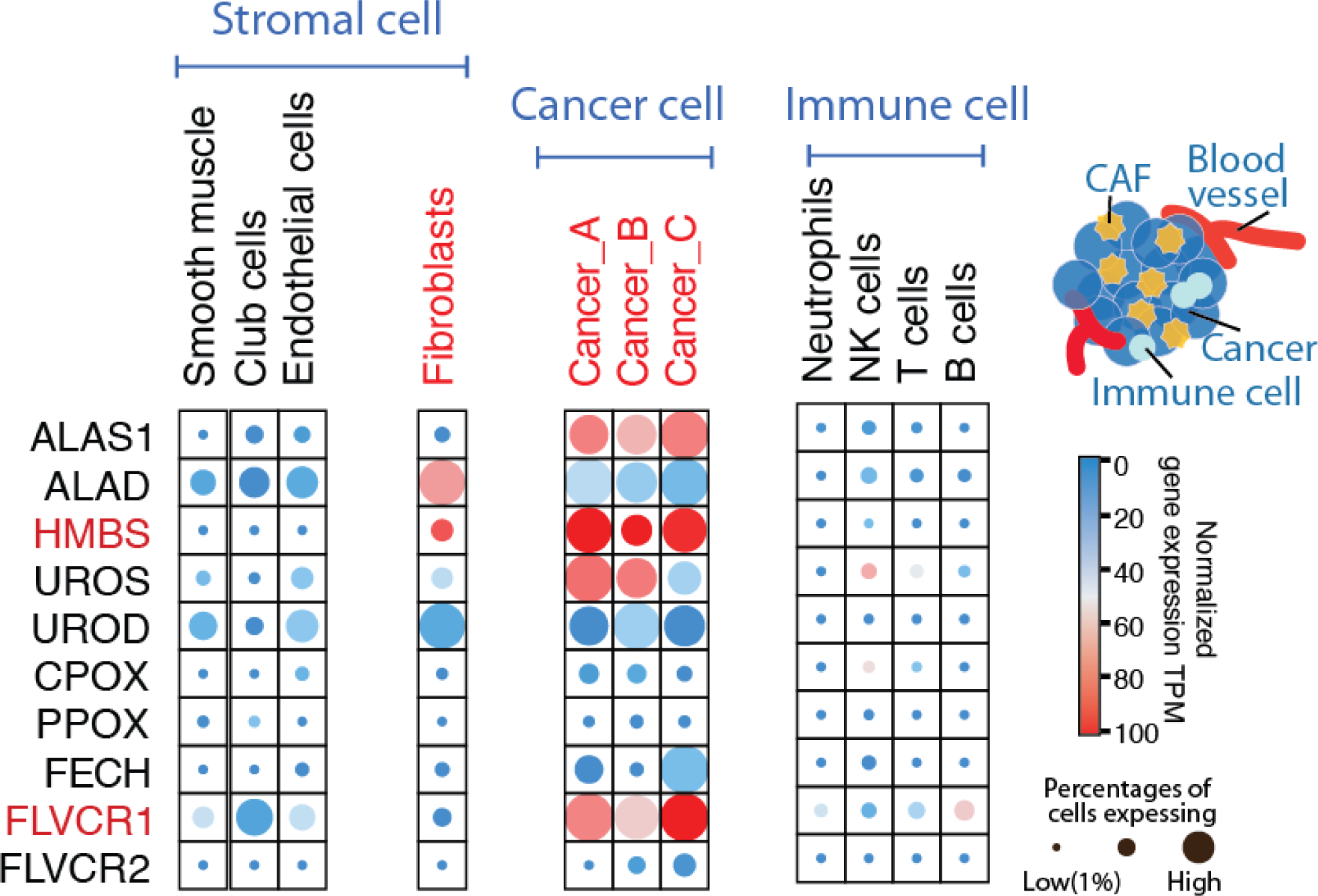
Single-cell RNA sequencing of cells from human lung cancer patients supports non-homeostatic porphyrin production in cancer associated fibroblasts. The data on single-cell lung cancer transcriptomes were retrieved from Zilionis *et al*. The expression of the genes for the heme biosynthetic pathway enzymes and heme trafficking proteins was analyzed in diverse cell populations present in the tumor. The expression results associated with cancer cells from 3 different patients (Cancer_A n = 203 cells, Cancer_B n = 766 cells, and Cancer_C n = 538 cells) are plotted – *HMBS* and *FLVCR1* show the most upregulated gene expression patterns. In the major groups of stromal and immune cell populations, most cell types have no evidence of porphyrin accumulation; while cancer-associated fibroblasts (n=585) show evidence of porphyrin production with upregulation of *HMBS (p <0.05)*. Both *HMBS* and *FLVCR1* are significantly different between tumor and stromal cell populations (p <0.05). Color shading indicates normalized gene expression levels per gene, in a given cell population; and circle sizes indicate the percentage of cells in that population with detectable single-cell transcriptomes.

In cancer, porphyrin production is likely fostered by the specific TME, including CAFs [64]. Our single-cell RNAseq analysis indicated that lung CAFs show features of porphyrin production with upregulated HMBS gene expression (Fig. 3). Next, we provide the experimental validation that porphyrin production operates in CAFs derived from primary tumors from a lung adenocarcinoma patient, as assessed by PPIX quantification. The tumor is derived from treatment-naïve patient samples (female, 55 years, TNM Staging: T2N0M0). The samples were obtained from surgically excised tissues, and the CAFs were isolated (BioIVT). Immunohistochemical staining for Vimentin revealed that over 95% of isolated cells within the tumor are positive for Vimentin, indicating a significant presence of mesenchymal characteristics. We show that CAFs from primary lung adenocarcinoma accumulate heme intermediates (*e.g.*, PPIX) upon induction with ALA, a precursor substrate of the heme biosynthetic pathway, but they lose this feature when the CAFs are passaged *in vitro* for over 15 generations (Fig. 4A). Interestingly, the fibroblasts adjacent to the tumor also show porphyrin overdrive, albeit to a lesser extent (“CAF vs. Other fibroblasts adjacent to tumor”) (Fig. 4B). These findings suggest that TME acts as a continuum of aberrant heme metabolism.

**Fig. 4.**
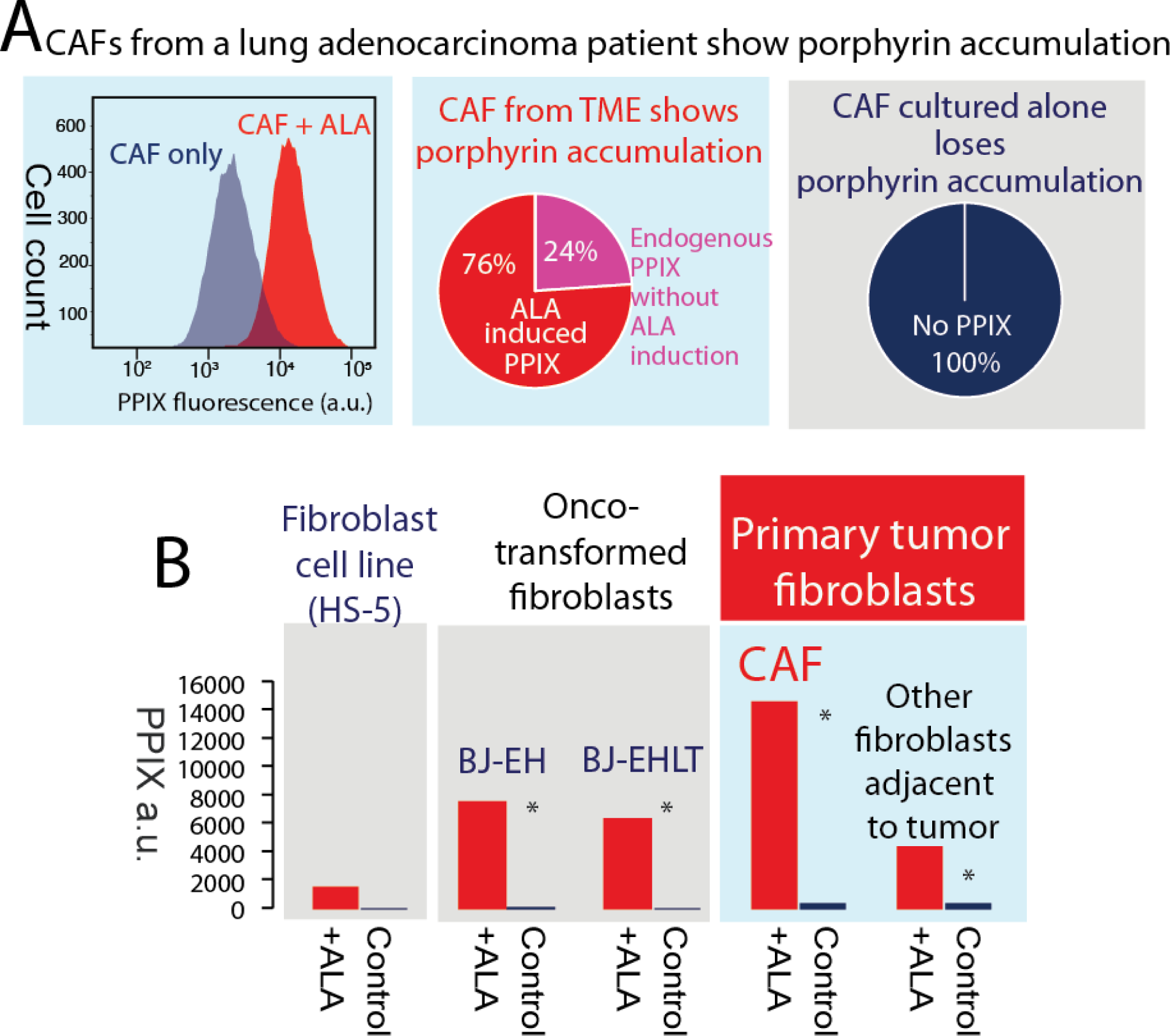
Primary CAFs accumulate PPIX. **A**. The primary tumor is primary lung adenocarcinoma. ALA (1 mM final concentration) was added at least 4 hours prior to flowcytometry and PPIX quantification. Normalized CAFs, after 15 *in vitro* passages from the same lung cancer patient, maintained rigorous cell growth and cell division but lost porphyrin production entirely (right panel). Porphyrin production, as estimated from PPIX accumulation, is present in over 99% of CAFs from primary lung adenocarcinoma (left panel). Even without ALA induction, PPIX accumulates in 24% of the CAFs (middle panel). **B.** Primary CAFs, onco-transformed fibroblasts and cancer cells show similar patterns of ALA induced PPIX. * Indicates significant differences.

In sum, our experiments show that the lung cancer TME exhibits characteristics of porphyrin production, based on single-cell RNAseq analysis and our experimental validation with CAFs isolated from a treatment-naive lung adenocarcinoma patient.

### Porphyrin overproduction is elevated in more aggressive cancers and linked to overall survival of patients

To evaluate the clinical relevance of porphyrin overdrive in lung cancers, we studied both patient tumor tissues from TCGA [45, 47] and analyzed previously generated single cell RNAseq data [65]. For human tissue gene expression in patient lung tumors, HMBS and FLVCR1 are the most enriched expressed genes, as compared to normal tissues (Fig. 5A). For the single cell transcriptomes, we used the single cell expression data from Zilionis *et al*. [65] and examined the expression differences in more aggressive vs. more differentiated lung cancer cell populations within the same tumor. Our results show both combined data of patient tumors (Fig. 5A), and single cancer cells show evidence of porphyrin production (Fig. 5B).

**Fig. 5.**
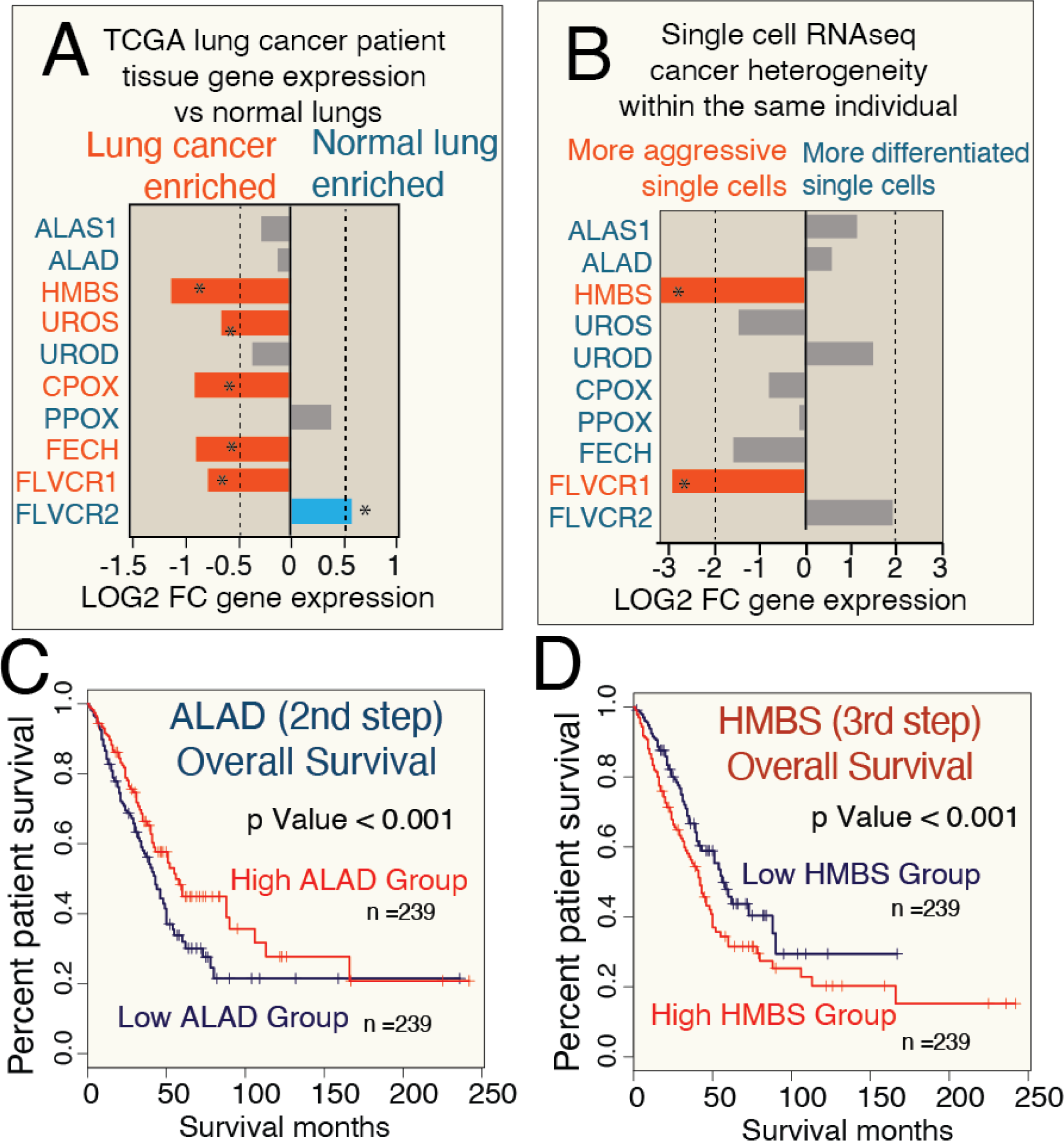
Lung cancer imbalanced heme biosynthesis is supported by both bulk and single cell RNAseq and is linked to the overall survival (OS) of patients. **A.** and **B.** Patient tumor tissue RNAseq and single cell lung cancer RNAseq support porphyrin overdrive. Genome analysis (shown in A.) was based on lung cancer data publicly available at The Cancer Genome Atlas (TGCA) and Genotype-Tissue Expression (GTEx) program. The single cell expression data from Zilionis *et al.* were used to study the expression differences in more aggressive *vs.* more differentiated lung cancer cell populations within the same tumor. **\*** denotes statistically significant differences (p <0.05) **C** and **D.** Imbalanced heme biosynthesis in lung cancer is related to patient OS. The Kaplan–Meier estimator, a non-parametric statistic, was used to estimate the survival function from the lifetime data. Shown are the Kaplan– Meier curves for data of visualization of *ALAD* (panel C) and *HMBS* expression (panel D) in lung adeno carcinoma cancer patients. The expression data were from TCGA and were analyzed in relation to the OS of the respective patients. Expression of *ALAD* and *HMBS* (the genes for the second and third enzymes of the heme biosynthetic pathway, respectively) show contrasting prognostic results in patients OS, suggesting that the greater the extent of porphyrin production (*i.e.,* the larger the difference between ALAD and HMBS expression), the more aggressive the lung cancer is. The statistical difference between the groups is estimated by the log-rank test. [FC, fold-change.]

We next analyzed overall survival with respect to high and low expression of the mid-step heme biosynthesis genes, ALAD and HMBS, using data from TCGA [47]. ALAD (the enzyme responsible for the 2nd step of heme biosynthesis) and HMBS (the enzyme that catalyzes the 3rd step) show contrasting prognostic results in overall survival, indicating the more imbalanced porphyrin production is (the larger the differences between ALAD and HMBS expression), the poorer the overall patient survival is (Fig. 5C,D).

## Discussions

TME is a dynamic milieu, orchestrating various cellular and molecular components. Dysregulated metabolisms within the TME have emerged as pivotal players in cancer progression, contributing to diagnostic nuances and therapeutic potential [66]. Metabolic shifts, such as the Warburg effect, heightened glutamine metabolism, altered lipid metabolism, perturbed one-carbon metabolism, and disrupted redox homeostasis, represent different aspects of metabolic rewiring in cancer [5, 6]. Our study introduces a novel facet of this metabolic landscape by delineating the dysregulation in heme biosynthesis, specifically the aberrant production of porphyrins, within the TME. The dysregulated heme metabolism, in form of porphyrin overproduction of both cancer cell and CAFs, suggesting a potential therapeutic vulnerabilities.

Notably, CAFs, recognized for their influential role in shaping the TME [6, 64], contribute to porphyrin production. The experimental validation of porphyrin accumulation in CAFs from primary lung adenocarcinoma emphasizes the contributions of different stromal players of aberrant heme metabolism within the TME. These findings suggest that the TME acts as a specialized microenvironment, fostering porphyrin production and influencing the overall metabolic landscape of cancer. The precise molecular pathways and trafficking routes of porphyrins within TME is yet to be determined.

The unexpected non-balanced reliance on intermediate steps of heme biosynthesis, as opposed to the initial and terminal steps, challenges conventional assumptions. We recently reported this process as ‘Porphyrin Overdrive’[38], and suggested it as a novel therapeutic target. Importantly, cancer cells exhibit a heightened dependence on mid-step enzyme genes, emphasizing the unbalanced nature of heme biosynthesis. This genetic vulnerability opens avenues for targeted therapeutic interventions. The significance of heme trafficking genes, particularly FLVCR2, offering potential targets for therapeutic strategies aimed at disrupting cancer-specific metabolic trafficking as a treatment to be developed.

Lung cancer is a pervasive global health concern, this type of cancer exhibits profound metabolic dysregulations that significantly impact its clinical outcomes [67, 68]. This study shows the absence of porphyrin accumulation in normal cells, in stark contrast to the presence of porphyrins in lung cancer cells. The evidence of porphyrin production in TME, as verified in lung adenocarcinoma, provides crucial insights into the metabolic rewiring of TME components associated with tumorigenesis. The clinical implications of porphyrin production in lung cancers extend beyond diagnostic considerations. The examination of patient tumor tissues and single-cell RNAseq data shows the association of porphyrin production with more aggressive cancers. High expression of mid-step heme biosynthesis genes correlates with overall poorer survival outcomes in lung cancer patients. This prognostic relevance establishes porphyrin production as a potential biomarker for predicting cancer aggressiveness and clinical progression. The integration of these findings into clinical practice holds the promise of refining prognostic assessments and tailoring therapeutic strategies based on the metabolic nuances observed in individuals cancer cases.

## Conclusions

Our investigation highlights a distinct facet of cancer metabolism characterized by the overproduction of porphyrins within tumors and their microenvironments. This aberrant porphyrin production, evident in both cancer cells and stromal components of the tumor microenvironment (TME), introduces a new dimension to the metabolic interplay of tumorigenesis. The exclusive presence of porphyrin overproduction in tumors and the TME, absent in normal tissues, demonstrates its potential as a novel therapeutic target. Additionally, porphyrin overproduction emerges as a promising prognostic indicator of cancer aggressiveness and clinical outcomes, suggesting avenues for innovative anti-tumor therapies aimed at disrupting metabolic interactions within the TME.

## Acknowledgments

R.H.Y.J., S.R.A, and G.C.F have received Florida Department of Health grant support 9BC14, to perform part of the research. R.H.Y.J. also received WLP (Women’s Leadership and Philanthropy) award support during research and writing of the work. Genetic onco-transformed fibroblasts were kindly provided by Robert Weinberg at MIT.

